# Evaluating Evolutionary and Gradient-Based Algorithms for Optimal Pathfinding

**DOI:** 10.1101/2025.03.16.643541

**Authors:** Olena Doroshenko

## Abstract

Pathfinding in complex topographies poses a challenge with applications extending from urban planning to autonomous navigation. While numerous algorithms offer potential solutions, their comparative efficiency and reliability when confronted with nonlinear terrains remain to be systematically evaluated. This study assesses three pathfinding algorithms—Genetic Algorithm (GA), Particle Swarm Optimization (PSO), and Sequential Quadratic Programming (SQP)—to establish a basis for comparison in terms of efficiency and computational speed. Results from twenty simulations indicate that SQP achieves lower path costs and reduced computational time than GA and PSO. In particular, SQP demonstrates reduced variability in path costs and quicker convergence to optimal paths, proving more effective in nonlinear environments. These results suggest gradient-based SQP as a preferable solution for complex pathfinding tasks. The study offers a framework for algorithm selection where efficiency and promptness are critical, potentially guiding decisions in operational strategies and system architecture.

## 1 Introduction

Modern logistics systems and decision-making processes in various industries use pathfinding algorithms to enhance efficiency and ensure safety. For example, in urban environments, these algorithms power traffic management systems that assess and modify vehicle routes, reducing travel delays and decreasing carbon emissions (Ghaffari et al. 2022; Boriboonsomsin et al. 2012; Namoun et al. 2021). In military operations, they help avoid resource-consuming or hazardous zones (Mora et al. 2013), adjusting routes based on threats or intelligence to protect troops and civilians. In emergency healthcare, pathfinding algorithms optimize ambulance routes by analyzing historical traffic and emergency data to predict and minimize delays when time is of essence (Michael and Xavier 2018; Mohd Nordin et al. 2011). Yet, while these and other (Murrieta-Mendoza, Botez, and Félix Patrón 2015; Wang, Liu, and Li 2023; Baker and Ayechew 2003) applications demonstrate the adaptability of pathfinding algorithms across various domains, determining the most effective algorithm for specific tasks remains an ongoing area of research.

Recently, three optimization algorithms such as Genetic Algorithm (GA), Particle Swarm Optimization (PSO), and Sequential Quadratic Programming (SQP) have shown considerable promise in solving pathfinding tasks. For instance, GAs have been used to optimize shelter locations and evacuation routes during emergencies (Yin, Zhao, and Lv 2023; Bayram 2016; Pourrahmani, Delavar, and Mostafavi 2015) by simulating a series of mutations and selections to iteratively evolve solutions toward the most efficient route under given constraints. Such genetic process regularly involves choosing the best outcomes as “parents” to produce “children” in the next generation, using selection based on performance scores, random crossover of genetic material, and mutations to maintain diversity (see De Jong 1988 for overview). However, while GAs are flexible and robust, offering high accuracy even for diverse environments, they can suffer from slow convergence rates and may get stuck in suboptimal solutions if not properly initialized (Vasconcelos et al. 2001). The second algorithm, PSO, inspired by the social behavior of animals (Kennedy and Eberhart 1995), proves beneficial for drone navigation, where multiple agents find optimal paths without centralized control (Mesquita and Gaspar 2021; Zhicai, Jiang, and Hong 2021; Kumar et al. 2022). Here, agents, or “particles,” scattered across a map, similarly to GA, adjust their paths based on the cost function values. Notably, while PSO is effective in converging to a solution, it too may get stuck in local optima. However, due to the exploration of the environment with multiple agents, such suboptimal outcome is less probable (Engelbrecht 2013). Finally, the third algorithm, SQP, has been applied in aerospace applications, where it was used to compute optimal flight trajectories that consider multiple objectives and constraints, such as fuel efficiency and time (Huang, Lu, and Nan 2012; also see Hammad et al. 2020 for review). While SQP excels in precision when handling diverse environments and (compared to the previous two algorithms) can provide solutions faster, like PSO and GA, it can struggle with local minima. Its deterministic nature and absence of evolutionary-inspired behavior may require modifications or enhancements to ensure broader exploration of the solution space.

In this study, I benchmarked three pathfinding algorithms in a complex nonlinear environment (Figure 1A) to identify the algorithm that offers optimal time efficiency and accuracy. Each algorithm underwent evaluation for simulation latencies and path costs, with further details on the cost function provided in the Methods section. I executed twenty simulations per algorithm, establishing statistical distributions for both cost and latency metrics. Through statistical comparison, I identified the most effective algorithm for this specific pathfinding task.

**Figure 1.**
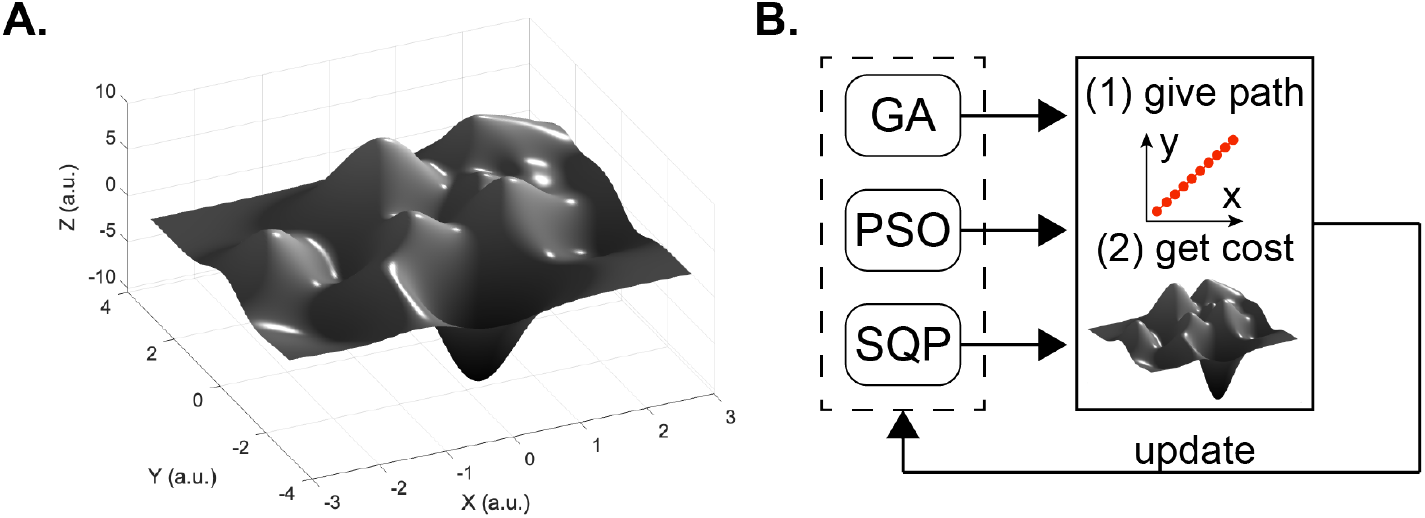
The environment to traverse (**A**) and the simulation pipeline (**B**). The environment was generated as a linear combination of scaled and translated Gaussian surfaces, as described in Methods and Materials. The simulation procedure entailed refining the path predictions by each algorithm separately using the landscape-dependent cost function. Abbreviations: GA—genetic Algorithm; PSO—Particle Swarm Optimization; SQP— Sequential Quadratic Programming; a.u.—arbitrary units.

## 2 Methods and Materials

### 2.1 Environment

The environment for the path optimization was defined as a linear combination of scaled and translated Gaussian surfaces generated using the “*peaks*” function (MATLAB, The MathWorks, Inc., Natick, MA, USA). The environment is depicted in Figure 1A, and its MATLAB code is available in the Supplementary Materials. The domain for path traversal was constrained within [−3, 3] arbitrary units (a.u.) on both *x* and *y* axes, with no constraints along the *z*-axis. The start and end points of the path were defined at [−2.8, −2.8] and [2.8, 2.8] a.u., respectively, to mitigate boundary condition impacts during traversal. A fixed number of ten waypoints, *P* = {*P*_1_, *P*_1_, … *P*_*n*_}, including the start and end points, was chosen to balance path fidelity and optimization time.

### 2.2 Cost Function

The cost function for traversing the environment, denoted *C*(*x, y*), was defined based on the absolute value of the *z*-coordinate at specific points along a defined path on the *x*— *y* plane. For cost evaluation, each segment between consecutive waypoints *P*_*i*_ and *P*_*i*+1_ was interpolated into smaller segments of 0.1 a.u. The cost for each interpolated point (*x, y*) was evaluated, summated, and multiplied by the distance to the subsequent interpolated point:

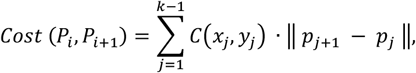

where *p*_*j*_ and *p*_*j*+1_ are the interpolated points between *P*_*i*_ and *P*_*i*+1_, and *k* = 10 is the number of interpolated points between these two path points.

The overall cost of traversing the complete path was determined by summing the costs of all individual segments, adjusted by a coefficient that incorporates the total length of the path to penalize longer routes:

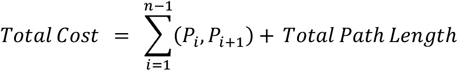

The last term in this expression aligned optimization objective with the practical requirement to minimize displacement along the *x*— *y* plane.

### 2.3 Optimization Algorithms

To find optimal path, as depicted in the simulation pipeline schematic in Figure 1B, I used three algorithms—two evolutionary and one gradient-based.

#### Genetic Algorithm

The first evolutionary algorithm applied was the Genetic Algorithm (GA), a search method that mimics the process of natural selection (see Mitchell 2001 for overview). The algorithm was initiated with a randomly generated population of solutions, each comprising ten waypoints *P* = {*P*_1_, *P*_1_, … *P*_*n*_} that defined the path. The population was set to 100 solutions—referred to as “*individual*s,”—to balance genetic diversity and simulation efficiency. The algorithm was configured to execute over a maximum of 100 generations, during which selection, crossover (crossover fraction was set to 0.8), and mutation (random mutation percentage was chosen from a Gaussian distribution) were applied to each generation to refine the population towards the optimal path. These operations introduced variability and prevented the algorithm from being trapped in local optima, increasing the likelihood of finding a global optimum.

#### Particle Swarm Optimization

Second evolutionary algorithm, Particle Swarm Optimization (PSO) (Kennedy and Eberhart 1995), a method inspired by the social behavior of mammals, relied on collective intelligence of individuals, referred to as “*particles*”, simultaneously exploring the environment to find minima. The movement of particles through the environment was described by their position *q* and velocity *ν* in the environment:

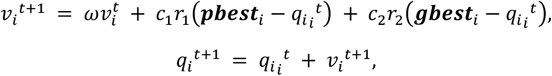

where *ω* is the inertia weight that controls the impact of the previous velocity on the current velocity; *c*_1_ and *c*_2_ are cognitive and social scaling coefficients, respectively; *r*_1_ and *r*_2_ are random numbers between 0 and 1; ***pbest***_*i*_ is the best position that particle *i* has visited; ***gbest***_*i*_ is the best position any particle in the swarm has visited.

To ensure parity in computational effort with the GA, I configured the PSO with a swarm of 100 particles and limited the optimization process to 95 iterations. This setup mirrored the population of the GA and resulted in similar number of function evaluations (9605 times for GA and 9600 times for PSO). Algorithmically, however, unlike the GA, which adhered to survival of the fittest, PSO used both the individual-best and the global-best solutions found by the swarm, which was expected to give it an advantage in escaping local minima.

#### Sequential Quadratic Programming

The third algorithm used in our study was a quasi-Newton method—the Sequential Quadratic Programming (SQP) procedure (“*fmincon*” function in the MATLAB’s Optimization Toolbox)—described in detail in (Boggs and Tolle 1995). Briefly, SQP is well-suited for constrained nonlinear optimization problems, where it operates by iteratively solving quadratic approximations to the Lagrangian of the cost function. Here, the algorithm was initiated with an initial guess defined as a straight path trajectory between start and end points. To this baseline, a white noise perturbation was introduced at each call to the algorithm. The noise was envisioned to aid SQP in escaping local minima and ensure exhaustive search across the set of solutions defined by the constraints: −3 ≤ *P*_*i*_ ≤ 3 for each of the coordinates of the ten waypoints. Again, to ensure fair comparison across the algorithms, the total number of function evaluations was capped at 9605, matching the evaluation number used by GA at 100 generations. This setting also aligned with that of Particle Swarm Optimization (PSO), which used a marginally lower count of evaluations at 95 iterations, differing by only five evaluations. Unlike GA and PSO, however, the SQP is particularly efficient at rapidly converging to a local minimum, especially when the initial guess is close to the optimal solution.

### 2.4 Statistical and Performance Analyses

The performance of each algorithm was assessed by evaluating (1) precision, measured by the cost of the computed path, and (2) simulation execution time, assessed over 20 simulation sessions. Distributions of path costs and execution times were tested for normality using the Kolmogorov-Smirnov test. Normal distributions were compared using a one-way analysis of variance (ANOVA) test. The significance level (α) was set at 0.05. All simulations and analyses were conducted using MATLAB (The MathWorks, Inc., Natick, MA, USA).

## 3 Results

The three algorithms were used to compute the optimal paths. Twenty simulations were conducted for each algorithm to generate distributions of predicted path costs and execution latencies—the time it took for the algorithms to compute the path. Representative paths computed by each algorithm are shown in Figure 2. As can be seen in the figure, each algorithm produced a qualitatively different trajectory, with GA (Figure 2A) and SQP (Figure 2C) producing more realistic solutions that traverse the environment as expected. In contrast, PSO takes a shortcut (Figure 2B). Whether this shortcut was an exploitation of the environment’s weaknesses, or a more cost-efficient strategy was determined from the statistical comparisons between the cost distributions computed by each algorithm.

**Figure 2.**
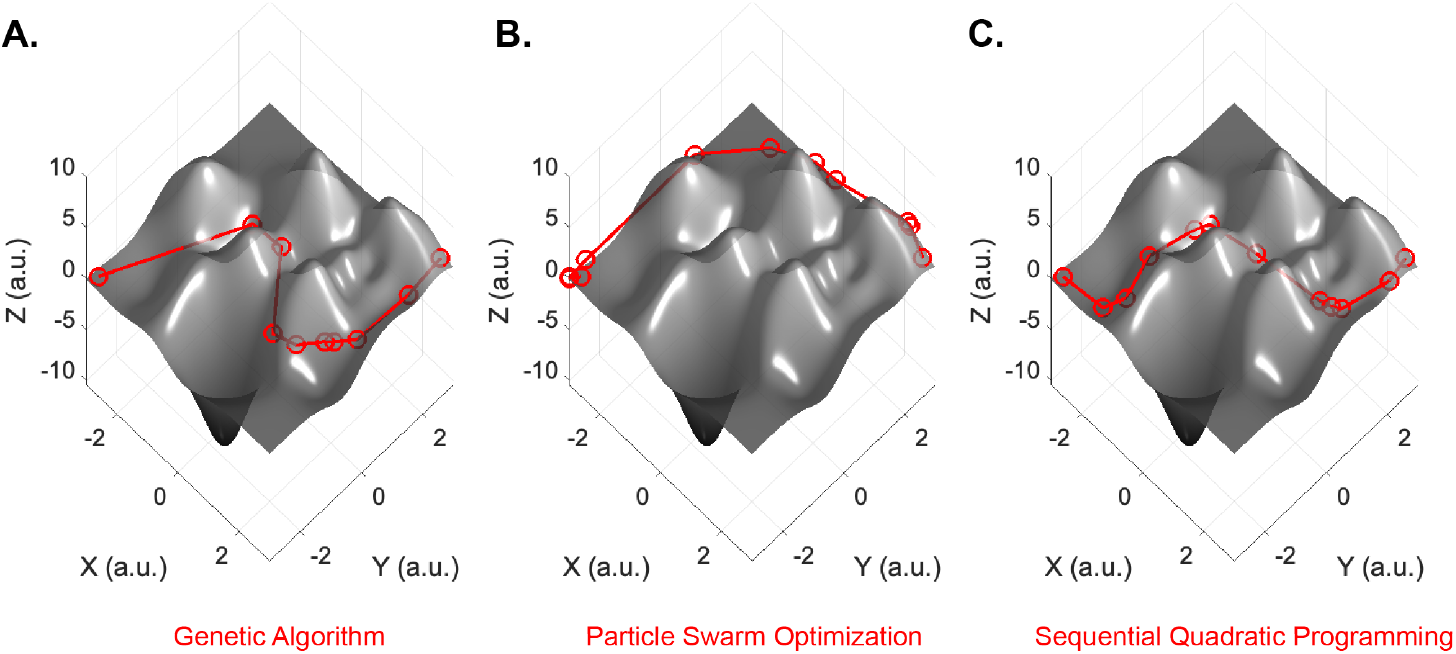
Representative paths calculated by the three evaluated algorithms: (**A**) Genetic Algorithm (GA), (**B**) Particle Swarm Optimization (PSO), and (**C**) Sequential Quadratic Programming (SQP), each illustrating the distinct pathfinding solutions. Abbreviations are the same as in Figure 1.

Figure 3A presents the comparisons between the final costs of the paths computed in each of the 20 simulations conducted by the three methods. All distributions were confirmed to be normal (Kolmogorov-Smirnov test, p < 0.05), and their differences were assessed using a one-way analysis of variance (ANOVA). The performance of the GA showed the greatest variability, with a standard deviation (STD) of 1.9381 a.u.; for PSO and SQP, the STDs were 0.9035 a.u. and 0.1199 a.u., respectively. High variability in path planning is undesirable as methods should ideally converge on the best solution in each simulation. However, of greater importance is the low cost of the solution. The ANOVA revealed no significant differences in the final costs between GA and PSO (p < 0.01), indicating that despite the greater variability in the distribution of solutions by GA, both evolutionary methods performed equivalently in terms of cost. This results also indicates that the map exploitation strategy used by PSO was not beneficial. In contrast, the performance of both evolutionary algorithms was significantly worse than that of the gradient-based SQP (p < 0.01), which demonstrated both lower variability in its predictions and robustness to noise corruption of initial conditions, highlighting its superior optimization capabilities.

**Figure 3.**
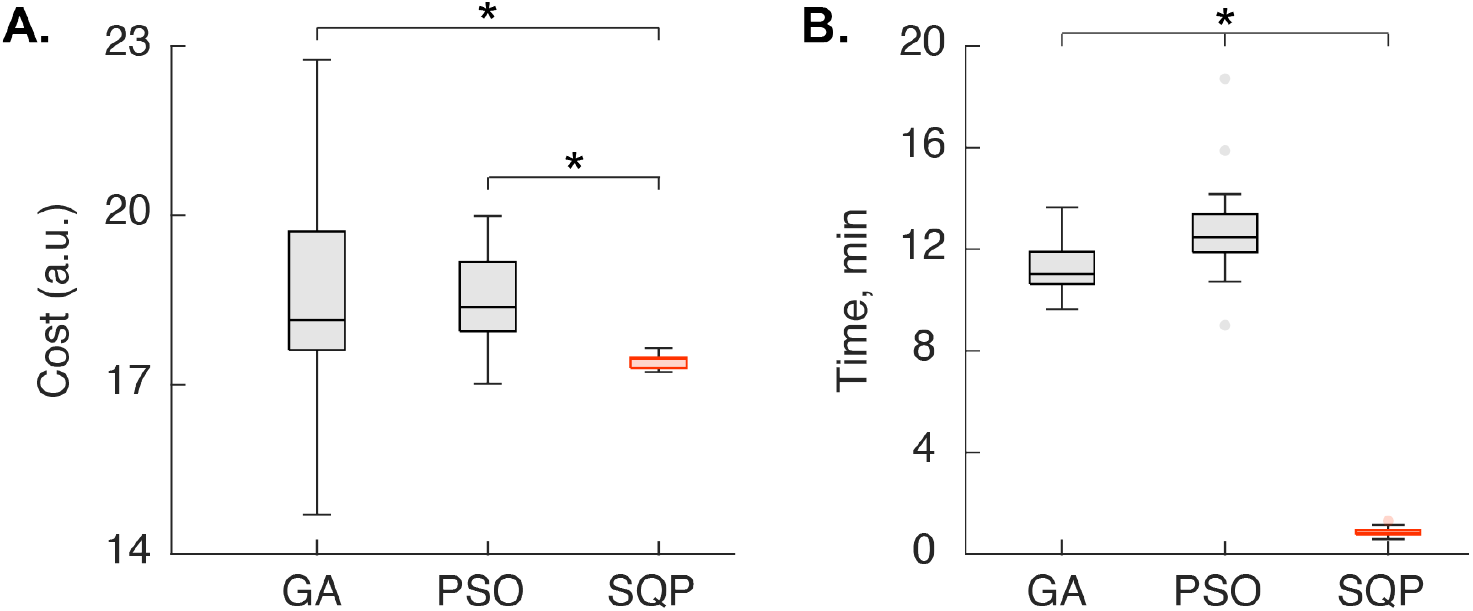
Distributions of path costs (**A**) and simulation execution times (**B**) for 20 simulations across three methods. The algorithm with superior performance is highlighted in red in each subplot. Asterisks denote statistically significant differences. Boxes show the interquartile range, bounded by the 25th and 75th percentiles, with whiskers extending to the non-outlier minimum and maximum values. Outliers are indicated as individual points. Abbreviations are the same as in Figure 1.

Figure 3B presents the distributions of simulation execution times. Similar to the analysis of cost distributions, these were tested for normality (Kolmogorov-Smirnov test, p < 0.05) and compared using ANOVA. The normally distributed execution times for all three methods differed significantly (p < 0.01). PSO demonstrated the longest execution time, followed closely by GA, while SQP was the fastest. The faster performance of SQP can be attributed to its gradient-based approach, which generally converges more quickly to a solution by following the steepest descent path. GA, which relies on a population-based search method, typically requires more iterations to converge, accounting for its slower speed relative to SQP but faster than PSO. PSO, while also a population-based method, often involves more complex calculations per iteration and may require additional adjustments to avoid local optima, leading to longer execution times.

## 4 Conclusions

This study evaluated the efficiency and performance of three pathfinding algorithms— Genetic Algorithms (GA), Particle Swarm Optimization (PSO), and Sequential Quadratic Programming (SQP)—through twenty simulations examining path costs and execution latencies. The findings revealed distinct behaviors and efficiencies: SQP consistently demonstrated superior performance with the lowest variability and fastest execution times, attributed to its gradient-based methodology that ensures quicker convergence. In contrast, GA and PSO showed higher variability in their path cost distributions, with PSO in particular showing the longest execution times due to its complex iterative calculations and potential to get stuck in local optima. Despite similar costs between GA and PSO, the significantly lower performance variability and greater robustness of SQP against initial condition perturbations underscore its advantages for complex pathfinding tasks in diverse environments. This study’s insights into the comparative efficiencies and behaviors of these algorithms provide valuable implications for selecting appropriate pathfinding techniques based on specific operational criteria and environmental contexts.

## Acknowledgements

I thank Serhii Bahdasariants for reviewing this work and providing detailed advice on its computational implementation during our meetings. I also thank Dr. Marjorie Darrah for her continued support of this project.

